# A semisynthetic glycoconjugate provides expanded cross-serotype protection against *Streptococcus pneumoniae*

**DOI:** 10.1101/2021.07.29.454378

**Authors:** Paulina Kaplonek, Ling Yao, Katrin Reppe, Franziska Voß, Thomas Kohler, Friederike Ebner, Alexander Schäfer, Ulrike Blohm, Patricia Priegue, Maria Bräutigam, Claney L. Pereira, Sharavathi G. Parameswarappa, Madhu Emmadi, Petra Ménová, Martin Witzenrath, Sven Hammerschmidt, Susanne Hartmann, Leif E. Sander, Peter H. Seeberger

**Author notes:** Co-senior authors, co-correspondence authors.

## Abstract

*Streptococcus pneumoniae* infections are the leading cause of child mortality globally. Current vaccines fail to induce a protective immune response towards a conserved part of the pathogen, resulting in new serotypes causing disease. Therefore, new vaccine strategies are urgently needed. Described is a two-pronged approach combining *S.pneumoniae* proteins, pneumolysin and PspA, with a precisely defined synthetic oligosaccharide, whereby the carrier protein acts as a serotype-independent antigen to provide additional protection. Proof of concept in mice and swine models revealed that the conjugates inhibit colonization of the nasopharynx, decrease the bacterial load and reduce disease severity in the bacteria challenged model. Immunization of piglets provided the first evidence for the immunogenicity and protective potential of synthetic glycoconjugate vaccine in a large animal model. A combination of synthetic oligosaccharides with proteins from the target pathogen opens the path to create broadly cross-protective (“universal“) pneumococcal vaccines.

## Introduction

Infections with *S. pneumoniae*, causing pneumonia, septicemia, and meningitis, are a leading cause of mortality globally [1], and pneumococcal diseases are a major cause of death in children under five years in developing countries [2]. According to the World Health Organization report from 2018, more children (800,000) under the age of five died from pneumococcal diseases worldwide than AIDS, malaria, and measles combined [3]. In the USA, *S. pneumoniae* infections are the most common reason for children to be hospitalized, and about one million adults seek hospital care annually [4, 5]. As the leading cause of bacterial meningitis in people over 65, *S. pneumoniae* causes substantial morbidity and mortality [5]. Considering the significant impact of pneumococcal diseases, surprisingly little is invested in vaccines and other preventive measures to address this problem. Currently marketed pneumococcal conjugate vaccines (PCVs) contain capsular polysaccharides (CPS) isolated from the serotypes associated with invasive pneumococcal diseases (IPDs). The 23-valent polysaccharide vaccine (Pneumovax 23^®^) is not effective in young children and the elderly [6, 7], while 7-, 10- and 13-valent polysaccharide-protein conjugate vaccines (Prevnar^®^ or Synflorix^®^) provide significantly increased protection in all age groups [8]. However, the manufacture of PCVs relies on the conceptionally simple but operationally challenging isolation of CPS from cultured bacteria. The presence of impurities, batch-to-batch variations, and unpredictable conjugation to a carrier protein can be overcome when synthetic oligosaccharides are employed. A rapidly growing field of medicinal chemistry allows the chemical synthesis of complex glycans on a large scale and improves the purity, homogeneity, and reproducibility of the antigen [9]. Synthetic oligosaccharides resembling the CPS of different serotypes can serve as candidate antigens for multiple vaccines and are a promising approach for the glycoconjugate vaccine development [10, 11].

The protection levels of marketed multivalent pneumococcal PCVs vary significantly depending on geographical location and serotype prevalence [12]. Colonizing serotypes covered by the vaccines are often replaced by previously less prevalent serotypes that acquired current colonizing strains under selective pressure [13]. The plasticity of the pneumococcal genome may allow for adaptation to the selective pressure of vaccines [14]. Thus, the development of a “universal” pneumococcal vaccine that is immunogenic in all age groups and provides broad serotype-independent protection against all serotypes has been pursued for decades [15]. Multivalent combinations of pneumococcal membrane proteins appear to afford additive protection against pneumococcal infection in mice [16].

*S. pneumoniae* proteins such as pneumolysin toxoid (Ply) [17, 18] and pneumococcal surface protein A (PspA) [19, 20] have been explored for cross-serotype protective pneumococcal vaccines. Ply is considered a major virulence factor of *S. pneumoniae* as it forms pores in the cell membrane that facilitate bacterial spread and increase barrier breakdown and host cell death, thereby contributing to disease manifestation [21, 22]. Analysis of mucosal immunity of children showed a significantly higher level of anti-Ply antibodies in children with negative pneumococcal culture compared to carriage-positive individuals, suggesting that anti-Ply antibodies are protective against colonization [23]. Animal models using Ply-deficient *S. pneumoniae* mutants have revealed decreased colonization of the nasopharynx, increased bacterial clearance from the lung, and prolonged survival following infection [24]. Pneumococcal surface protein A (PspA) is an essential surface molecule of *S.pneumoniae* which leads to inhibition of the complement-mediated bacterial clearance [25, 26]. The multifunctional immune evasive properties of PspA are essential for pneumococcal nasopharyngeal colonization and invasion of *S. pneumoniae.* PspA is immunogenic and induces both humoral and cellular immune responses [27]. Active immunization with purified PspA protects against nasopharyngeal carriage and invasive disease in animal models [16].

Serotype replacement following PCV introduction spurred research into broadly cross-protective (“universal“) pneumococcal vaccines. Glycoconjugate vaccines induce, in addition to the response to the glycan antigens, a substantial immune response against the carrier protein as an “additional valency”. Here, we explore the use of cross-protective conserved pneumococcal proteins as carriers for synthetic serotype-specific oligosaccharide antigens to broaden vaccine specificity and prevent serotype replacement. A detoxified version of Ply or PspA served as carriers for synthetic oligosaccharide antigens specific for *S. pneumoniae* serotypes 2 or 3. The carbohydrate- and protein-specific immunogenicity was robust, with potent opsonophagocytic and cross-neutralizing serum activity. These novel semisynthetic pneumococcal glycoconjugate vaccine candidates were protective against pneumococcal pneumonia and provided cross-protection against nasopharyngeal colonization with a heterologous serotype in mice. These are the first examples of glycoconjugates combining immunogenic pneumococcal protein carriers with defined synthetic oligosaccharide antigens demonstrating a promising strategy to improve protection against pneumococcal diseases.

## Methods

### Animal experiments

Animals were treated according to German (Tierschutz-Versuchstierverordnung) and European Regulations (Directive 2010/63/EU). Recommendations of the Society for Laboratory Animal Science (GV-SOLAS) and of the Federation of European Laboratory Animal Science Associations (FELASA) were followed. The permits used in the study were approved by the Office for Health and Social Affairs Berlin (LAGeSo); Permit Number: G0135/14, G0053/18; and the Landesamt für Landwirtschaft, Lebensmittelsicherheit und Fischerei Mecklenburg-Vorpommern (LALLF M-V, Rostock, Germany); Permit Numbers: 7221.3-1-061/17 and 7221.3-2-042/17. All efforts were made to minimize the discomfort of the animals and ensure the highest ethical standards.

### General Methods

Synthetic oligosaccharide S. pneumoniae serotype 3 tetrasaccharide (ST3) [28] and serotype 2 hexasaccharide (ST2) [29] were conjugated to carrier proteins CRM197 (EirGenix, Inc., Taiwan), purified Pneumolysin PlyW433E and PspA using the standardized protocol described previously [28, 29]. Synthetic antigens were printed on NHS-activated microarray slides. Mice and swine were immunized according to the prime + boost + boost schedule and infected with S. pneumoniae serotype 3 (PN36) and S. pneumoniae serotype 2 (D39. The immunogenicity of the vaccines was analyzed by glycan array and ELISA. The protective ability of the antibodies to trigger opsonophagocytic killing of bacteria and toxin neutralization were assessed by *in vitro* opsonophagocytic killing assay (OPKA), red blood cells lysis assay and electric cell-substrate impedance sensing (ECIS). The presence of carbohydrate specific T cells was analyzed by antigen-reactive T cell staining of pigs by flow cytometry. Detailed materials and methods can be found in SI Appendix.

## Results

### ST3-PLY glycoconjugate is immunogenic and protects against S. pneumoniae serotype 3 induced pneumonia

It has been shown that synthetic glycoconjugate vaccine candidate resembling the ST3 CPS repeating units protects mice against ST3 induced pneumonia [28]. Detoxified diphtheria toxoid (CRM197) is used as a carrier for CPS in PCV13. Here, we conjugated the ST3 tetrasaccharide to an immunogenic detoxified Pneumolysin (PlyW433E) [30] using a bifunctional linker (**Fig. 1a**). The ST3-tetrasaccharide Ply conjugate (ST3-Ply) contained on average seven synthetic tetrasaccharide molecules attached to each Ply protein (**Sup Fig. 1**).

**Figure 1.**
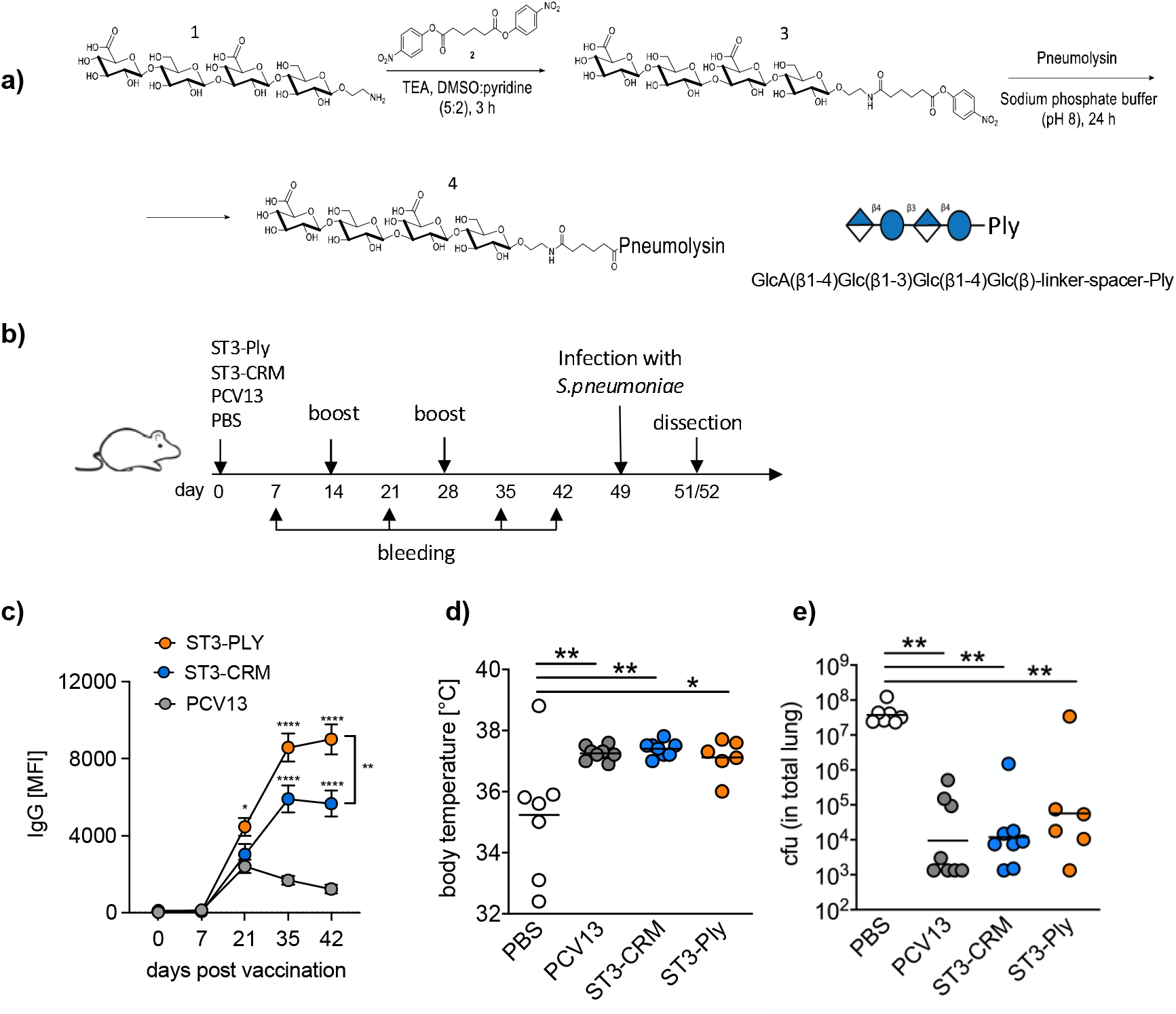
ST3-Ply glycoconjugate is immunogenic and protects against *S. pneumoniae* serotype induced pneumonia. **a)** ST3-tetrasaccharide and Ply conjugation reaction. Synthetic ST3-tetrasaccharide **1** was covalently conjugated with Ply carrier protein using *p*-nitrophenyl adipate ester **2** as a coupling reagent, yielding semisynthetic glycoconjugate ST3-Ply **4**. **b)** Vaccination regimen. ST3-Ply, ST3-CRM, and PCV13 were administered at indicated time points, followed by challenge with 5 × 10^6^ CFU of *S. pneumoniae* serotype 3 bacteria. Mice (*n* = 8) were immunized with a dose of vaccine equivalent to 0.4 μg of synthetic antigen. **c)** Kinetics of ST3-tetrasaccharide specific IgG measured by glycan array. Serum samples from mice immunized with ST3-Ply, ST3-CRM197, and commercial PCV13 vaccine were collected on days 0, 7, 21, 35, and 42. Dots indicate the mean fluorescence intensity value ± SEM of eight animals per group. Statistical analysis was performed by Tukey’s multiple comparisons test. **d)** Body temperature changes 48h after the pneumococcal challenge. Statistical analysis was performed by Tukey’s multiple comparisons test. **e)** Bacterial burden in lungs 48h post-infection. CFUs were calculated for each animal and plotted as individual data points from *n* = 8 mice (*n* = 6 for ST3-Ply vaccinated group). Statistical analysis was performed by Tukey’s multiple comparisons test. Statistical significance for all panels * p <0.05, ** p <0.01, *** p <0.001**** p <0.0001.

To assess the immunogenicity of the glycoconjugates, mice were immunized with alum-adjuvanted ST3-Ply and compared to ST3-CRM197 and PCV13 in a prime-boost regimen **(Fig. 1b)**. We observed significantly higher antigen-specific antibody levels for ST3-Ply and ST3-CRM197, starting at 21 days post-immunization compared to PCV13 (**Fig. 1c)**. ST3-Ply conjugate induced a more robust ST3-specific antibody response compared to ST3-CRM197. Therefore, the conjugation to alternative carrier protein increases the immunogenicity of carbohydrate antigen compare to traditionally used CRM197. Animals immunized with CPS containing Prevnar13^®^ showed mild cross-reactivity to the ST3-tetrasaccharide (**Fig. 1c**).

Next, ST3-Ply immunized animals were intranasally infected with 5 × 10^6^ CFU of *S. pneumoniae* serotype 3, while negative controls received PBS only. Clinical signs of disease, body weight, and body temperature were monitored every 12 hours following a clinical score. Bacterial outgrowth in the lung was examined 48 hours post-infection. ST3-tetrasaccharide and carrier protein-specific antibodies were analyzed by glycan microarray. Immunization with ST3-Ply as well as ST3-CRM197 and PCV13 significantly alleviated disease severity, as evidenced by the absence of hypothermia, compared to PBS control (**Fig. 1d**). The absence of clinical manifestations of disease correlated with a strongly decreased bacterial burden in the lungs (**Fig. 1e**). Hence, a semisynthetic ST3-Ply conjugate is immunogenic and is highly protective against serotype-homologous pneumonia.

### Anti-Ply antibody responses show cross-neutralization potential

In line with the protective effects against the pneumococcal challenge, immune-sera from mice vaccinated with ST3-Ply showed *in-vitro* protective activity measured by OPKA (**Fig. 2a**). No direct opsonophagocytic activity towards unrelated serotypes was observed, indicating that CPS-specific antibodies are required for binding and activation of complement in this assay. To further analyze the neutralizing effects of anti-Ply antibodies, post-immunization seru samples (day 35) were incubated with native Ply in the presence of human red blood cells (hRBCs). The hemoglobin release through erythrocyte lysis was measured at the absorbance *A*_540nm_ as an indicator of pneumolysin activity. Antibodies from ST3-Ply immunized mice exhibited potent toxin neutralizing activity blocking hRBC lysis, compared to pre-immune serum. The neutralizing effect was preserved up to a titer of 1:100 (**Fig. 2b**). Next, hRBCs were incubated with bacterial lysate of *S. pneumoniae* serotypes 3 and heterologous serotypes 2 and 8, and post-immunization serum. A significant reduction of hemoglobin released independently of the pneumococcal serotype was observed (**Fig. 2c**). The most significant reduction in cytotoxicity was seen with lysate of ST3, suggesting an additional neutralizing effect of specific anti-glycan antibodies. Residual hemolysis was the result of different toxins and lytic components present in the bacterial cell lysates.

**Figure 2.**
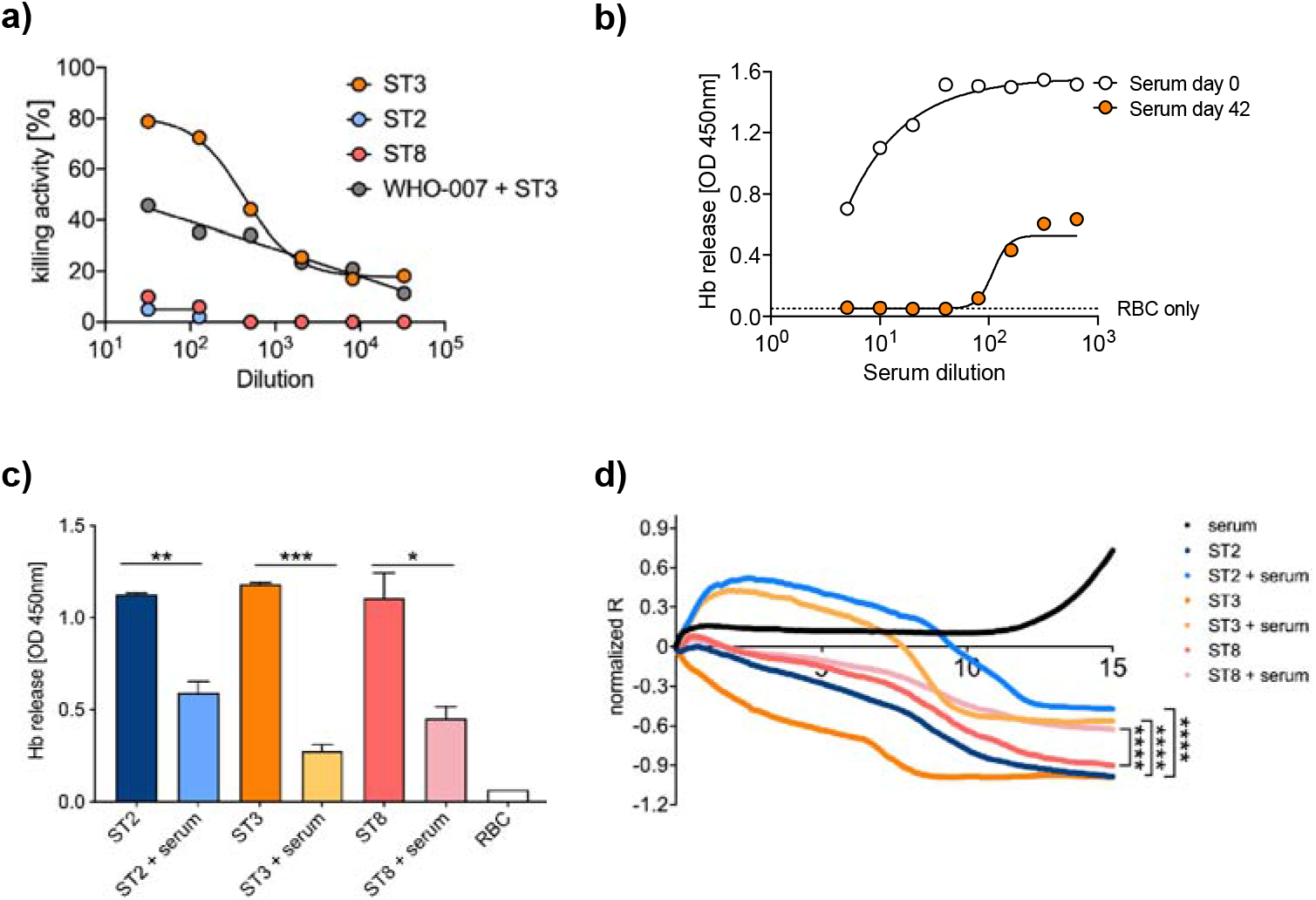
“*Ex vivo*” cross-protection and inhibition of serotype independent pneumolysin induced lysis of human red blood cells by anti-pneumolysin antibodies from mice immunized with ST3-Ply conjugate. **a)** The *in vitro* opsonophagocytic killing of *S. pneumoniae* ST2, ST3, and ST8 mediated by polyclonal antibodies from mice immunized with ST3-Ply glycoconjugate. Blood collected on day 35 after the immunization was used for the assay (*n* = 8). Data are means of CFU reduction relative to negative control wells (samples lacking either antibody or complement) of two independent experiments. WHO-007sp human control sera from volunteers immunized with pneumococcal polysaccharide vaccine were used as a standard. **b)** Inhibition of hemolytic activities of native pneumolysin (12.5μg) by antibodies from mice immunized with ST3-Ply conjugate. Different dilutions of serum from blood collected on days 0 and 42 were tested. Hemoglobin release is measured as the 540 nm absorbance (*A*_540_) of assay supernatants. **c)** The inhibition of hemolytic activities of whole-cell lysates of *S. pneumoniae* ST2, ST3, and ST8 by antibodies from mice immunized with ST3-Ply conjugate. Hemoglobin release is expressed as the *A*_540_ of assay supernatants (mean ± SD). The statistical significance was calculated by paired t-test. **d)** Serum from mice immunized with ST3-Ply conjugate protects pneumolysin-induced epithelial barrier disruption (two independent experiments). A549 cells were treated with the whole cell lysate of *S.pneumoniae* serotypes 2, 3, and 8 with and without serum from mice immunized with ST3-Ply (blood collected on day 35). Transcellular electrical resistance was continuously monitored for 15 hours and normalized to baseline resistance. Statistical significance was calculated by Wilcoxon matched-pairs signed rank test between bacteria lysate with and without antibodies. Statistical significance for all panels * p <0.05, ** p <0.01, *** p <0.001**** p <0.0001.

As epithelial cells of the respiratory tract are a primary target for *S. pneumoniae*, we analyzed the protective potential of vaccine-induced antibodies on maintaining barrier function *in vitro*. The human alveolar basal epithelial cells A549 were exposed to cell lysate of *S. pneumoniae* serotype 2, 3, and 8. Changes in transcellular electrical resistance of cell monolayers were analyzed by Electric Cell-substrate Impedance Sensing (ECIS). ST3-Ply post-immunization antibodies were able to significantly inhibit barrier breakdown measured as a loss of electrical resistance (**Fig. 2d**).

These results indicate that the ST3-Ply conjugate immunization triggers high levels of pneumolysin-neutralizing antibodies that may block acute lung injury in the early course of pneumococcal infection. Anti-toxin immunity, in addition to CPS-specific antibody responses, may exhibit additional benefits in the prevention of severe pneumococcal pneumonia.

### Intranasal immunization with ST3-Ply and ST3-PspA elicits mucosal and systemic immune responses and inhibits serotype-homologous and -heterologous colonization

*S. pneumoniae* virulence factors, such as Ply and PspA, play a crucial role in the initial step and the subsequent invasion and spreading of pneumococci [24]. Mucosal vaccine delivery may thus activate the local immune system against mucosal transmission and bacteria spread. Synthetic ST3-tetrasaccharide and ST2-hexasaccharide were conjugated to Ply and PspA to evaluate the benefits of those proteins as carriers and additional antigens for glycoconjugate vaccines. Glycoconjugates were characterized by MALDI-TOF-MS (**Sup. Fig. 2**).

Mice were immunized intranasally with ST3-Ply, ST3-PspA, ST2-Ply, ST2-PspA, or a mixture of both glycoconjugates according to the prime-boost regimen (**Fig. 3a**). Animals received a dose of glycoconjugate equal to 2.5 μg protein, corresponding to 0.5 μg synthetic oligosaccharide antigen, formulated with cholera toxin subunit B (CTB) as a mucosal adjuvant. Intranasal immunization triggered a systemic immune response in all mice tested. Both ST3-tetrasaccharide and CPS, as well as Ply- and PspA- specific antibodies, were induced in all mice receiving the vaccine with corresponding protein. (**Fig. 3b**).

**Figure 3.**
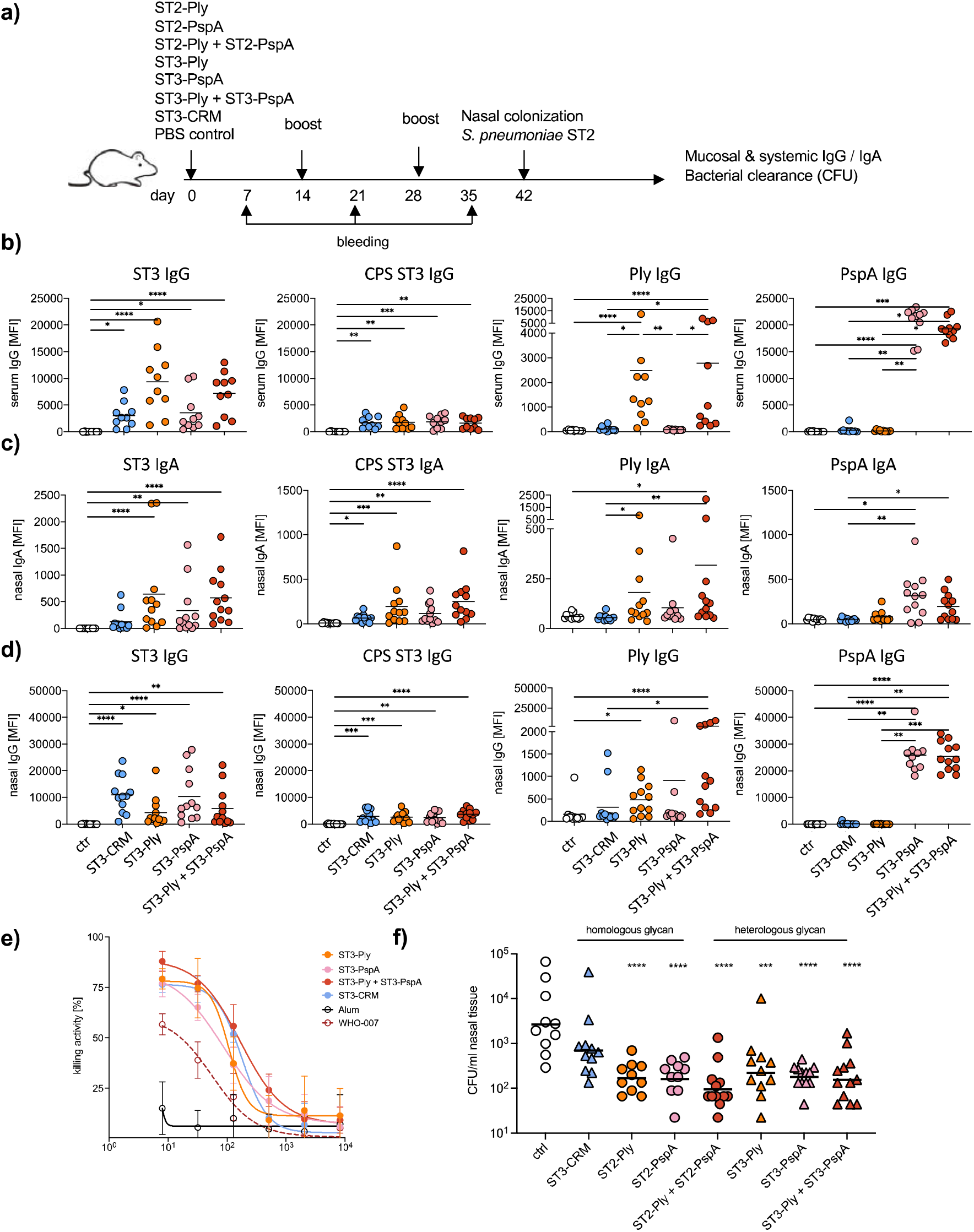
Intranasal immunization with ST3-Ply and ST3-PspA elicits mucosal and systemic immune responses and inhibits serotype-homologous and -heterologous colonization. **a)** Schematic overview of the study design. **b)** Serum IgG responses on day 35 post-immunization measured by glycan array expressed as MFI. Each dot represents an individual mouse (randomly chosen animals (*n* = 10) from each group). Statistical analysis was performed by Tukey’s multiple comparisons test. **c-d)** Evaluation of mucosal IgA (c) and IgG (d) responses on day 3 post intranasal immunization with the indicated conjugates to the indicated antigens (ST3-tetrasaccharide (ST3); CPS ST3; Pneumolysin (Ply) and PspA) measured in homogenized nasal tissue by glycan array expressed as MFI. Each dot represents an individual mouse (*n* = 12). Statistical analysis was performed by Tukey’s multiple comparisons tests. **e)** opsonophagocytic killing of *S.pneumoniae* serotype 3 mediated by antibody from mice immunized with ST3-Ply, ST3-PspA, and ST3-Ply + ST3-PspA glycoconjugates. Blood collected on day 35 was used for the assay. Data are means ± SD of CFU reduction relative to negative control wells (samples lacking either antibody or complement) of two independent experiments. **f)** Nasal colonization with *S. pneumoniae* serotype 2 (D39) three days after intranasal challenge of vaccinated C57BL/6N mice (*n* = 12) with 3.5 × 10^6^ CFU. Each dots represent CFU recovered from an individual animal. Data were analyzed using a one-way ANOVA multi comparison test. Statistical significance for all panels * p <0.05, ** p <0.01, *** p <0.001**** p <0.0001.

Two weeks after the final vaccination, mice were intranasally infected with a non-lethal dose of *S. pneumoniae* serotype 2. A local immune response (IgA and IgG) in homogenized nasal tissue was detected toward corresponding antigens. Mice injected with ST3-tetrasaccharide conjugates exhibited high antigen-specific antibody level respectively to the vaccine received (**Fig. 3c-d**). Animals immunized with ST2-hexasaccharide conjugates show similar mucosal IgA and IgG responses (**Sup. Fig.3**).

The protective activities of vaccine-induced antibodies were evaluated in post-immunization blood. ST3 conjugated vaccines were able to trigger the *in vitro* killing of *S. pneumonia* ST3 with efficiency higher than 75% (**Fig. 3e**). Intranasal vaccination with ST2-Ply, ST2-PspA, and ST2-Ply + ST2-PspA mixture, as well as an ST3-Ply, ST3-PspA, ST3-Ply + ST3-PspA mixture induced a significant reduction of bacterial load in the nasal cavity compared to PBS group (**Fig. 3f**). In summary, the vaccine-induced antibodies are able to mediate opsonophagocytic killing of bacteria and reduce the colonization as well as the risk of severe infection.

### Immunogenicity of ST3-CRM197 and ST3-Ply in Swine

Six-week-old healthy, female domestic pigs (German landrace *Sus scrofa*) were immunized intramuscularly with 2.2 μg of ST3-tetrasaccharide CRM197- or Ply-conjugated, dose equal to the PCV13, formulated with aluminium hydroxide. The control groups received PCV13 or Alum only. Blood for antibody responses was collected on days 0, 10, 20, and 35/36 post-immunization (**Fig. 4a**). The presence of ST3-tetrasaccharide and CPS-specific IgG_1_ and IgG_2_, as well as anti-carrier protein antibody, was analyzed by glycan arrays. CRM197 specific IgG_1,_ and IgG_2_ were identified in groups immunized with PCV13 and ST3-CRM197, whereas anti-Ply antibodies were mostly elicited by ST3-Ply vaccination (**Fig. 4b-e**). In groups vaccinated with semisynthetic ST3-CRM197 and ST3-Ply conjugates, ST3-tetrasaccharide specific IgG_1_ titers were significantly higher than IgG_2_ (**Fig. 4b**). A detectable level of a specific protein response was observed in unrelated groups due to nonsterile housing and breeding of swine. The study provides the first evidence for the immunogenicity of semisynthetic glycoconjugates in swine.

**Figure 4.**
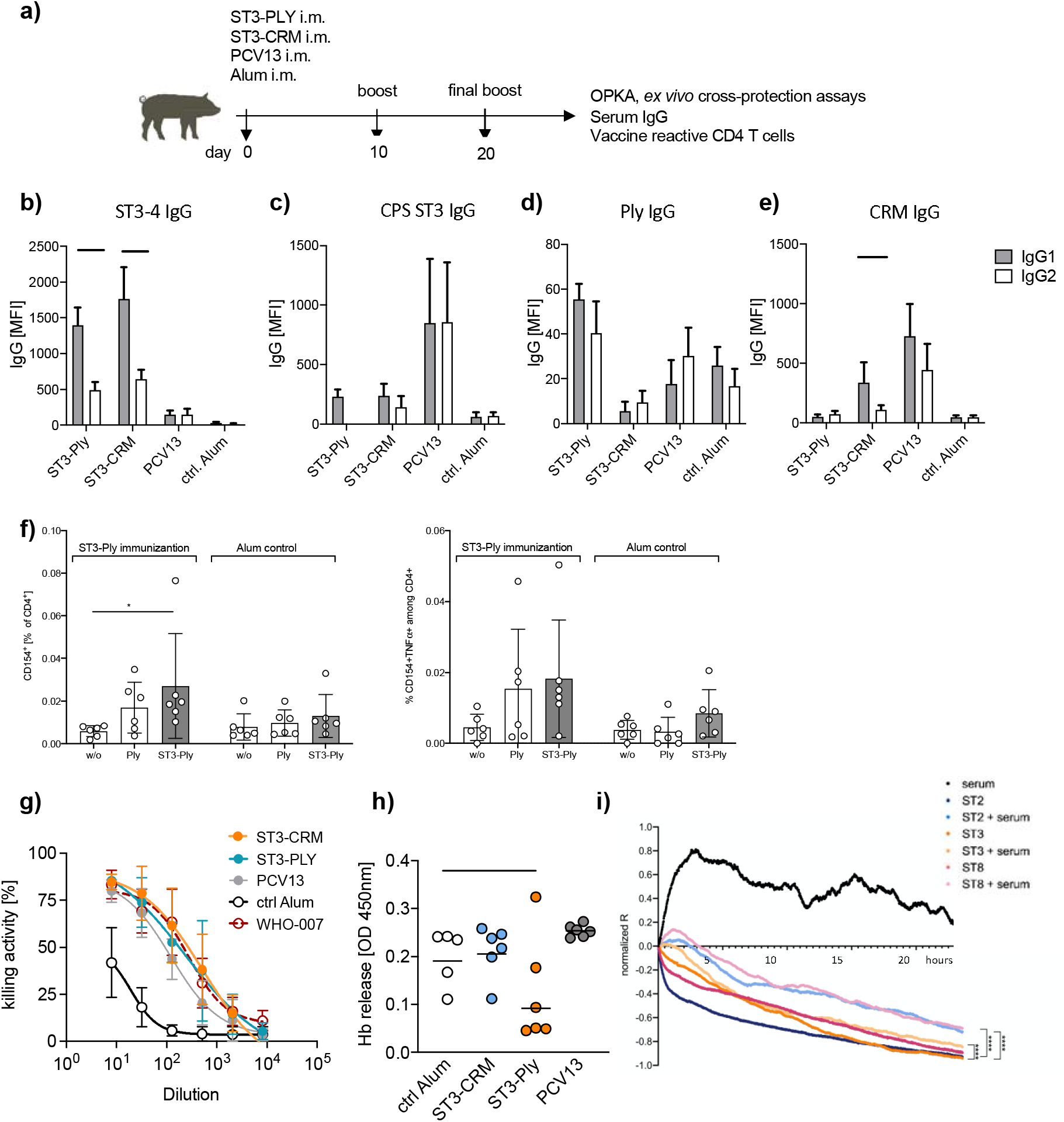
ST3-CRM197 and ST3-Ply glycoconjugates are immunogenic in swine. **a)** Schematic overview of the study design. **b-e).** Evaluation of IgG1/IgG2 responses in swine immunized with ST3-Ply or ST3-CRM197 or PCV13 and Alum adjuvant negative control. Antibody level against ST3-tetrasaccharide (b) and native *S. pneumoniae* serotype 3 CPS (c) and carrier proteins PLY (d) and CRM197 (e) measured by glycan array. Bars indicate the mean fluorescence intensity of each group ± SEM (*n* = 6 animals per group). Statistical significance was calculated by Wilcoxon Signed Rank test. **f)** Vaccine reactive CD4^+^ T cell responses measured following protein antigen re-stimulation for 6h of <PBMC and subsequent flow cytometry for total antigen-reactive CD4^+^CD154^+^ (left panel) or TNF-secreting CD4^+^CD154^+^ T cells (right panel). **g)** OPKA activity of pooled sera from pigs immunized as indicated. Graphs display means ± SD of CFU reduction relative to negative control wells (samples lacking either antibody or complement) of two independent experiments. Data were analyzed using a one-way ANOVA multi comparison test. **h)** Comparison of the toxin neutralization activities of immune sera (1:5 dilution) by measuring Hb release from sheep red blood cells, expressed as the A_540_ of assay supernatants. Each dot represents serum from one animal (*n* = 6). **i)** Serum from pigs immunized with ST3-Ply conjugate protects against pneumolysin-induced epithelial barrier disruption. A549 cells were treated with whole-cell lysates of *S. pneumoniae* serotypes 2, 3, and 8 with and without serum from pigs immunized with ST3-Ply (blood collected on day 20). Transcellular electrical resistance was continuously monitored and normalized to baseline resistance. Statistical significance was calculated by Wilcoxon matched-pairs signed rank test between bacteria lysate with and without antibodies. Statistical significance for all panels * p <0.05, ** p <0.01, *** p <0.001**** p <0.0001.

Next, we assessed antigen-specific T cell responses. We determined CD154 (CD40L) expression on CD4^+^ T cells restimulated with Ply or entire glycoconjugates to detect antigen-reactive CD4^+^ Th cells [31]. The frequency of antigen-reactive CD4 T cells in ST3-Ply immunized pigs was enhanced when compared to control animals (**Fig. 4f**). In response to ST3-Ply restimulation, a tendency towards an increased frequency of CD154^+^TNF^+^ CD4^+^ T cells was observed (**Fig. 4f**).

To further determine the protective capacity of ST3-tetrasaccharide specific antibodies, we performed OPKA using *S. pneumoniae* serotype 3. The serum from animals immunized with ST3-CRM197 and ST3-Ply showed similar OPKA activity to the PCV13 group (**Fig. 4g**). Thus, semisynthetic glycoconjugates ST3-CRM197 and ST3-Ply raise protective antibodies with strong opsonophagocytic activity in swine.

The post-immune sera from swine were evaluated for their toxin-neutralizing effect. Consistently, antibodies from pigs immunized with ST3-Ply potently inhibited pneumolysin-induced hemolysis of red blood cells *in vitro* compared to negative control and PCV13 groups (**Fig. 4h**). Since the animals were kept under normal husbandry conditions, some animals acquired neutralizing antibodies, probably following contact with pneumolysin-producing bacteria or cross-reactivity with other toxins. Nevertheless, the vaccine group immunized with ST3-Ply showed significantly decreased hemoglobin release compared to PCV13 vaccinated animals (*p = 0.0185*). Antibodies induced by ST3-Ply conjugate were able to inhibit the pneumolysin-mediated breakdown of barrier function in human alveolar basal epithelial cell line A549 *in vitro.* The transcellular electrical resistance of cell monolayers measured by ECIS showed significant differences while incubating with and without sera **(Fig. 4i).**

## Discussion

Bacterial pathogens that exist as a variety of different serotypes are a major challenge for the development of anti-bacterial vaccines. Multivalent vaccines, such Prevnar13^®^, cover only some of the 98 different strains of *S. pneumoniae,* which has allowed and driven replacement with non-vaccine serotypes (“serotype replacement“). The inclusion of conserved pneumococcal membrane proteins as vaccine antigens to create a “universal” *S. pneumoniae* vaccine has been investigated extensively [15, 32]. However, most bacterial proteins fail to provide sufficient protection since the bacterial capsule composed of CPS shields most membrane antigens and is the immunodominant target of anti-pneumococcal immune responses. In order to minimize serotype replacement while maintaining high levels of protection against IPD, we conjugated serotype-specific minimal synthetic CPS epitopes to immunogenic bacterial proteins of *S. pneumoniae* with the goal of combining the superior immunogenicity of CPS epitopes with the broader cross-serotype immunity of protein antigens.

A critical step in the development of pneumococcal infections is asymptomatic nasopharyngeal carriage [33]. Up to 10% of the adolescent population and more than 40% of young children, particularly in daycare, are colonized with *S. pneumoniae* [34]. Protective immunity against common serotypes of *S. pneumoniae* is required to protect risk groups against IPD. Therefore, vaccination strategies to target pneumococcal colonization need to target the mucosal immune response. Mucosal vaccines can induce both local and systemic immunity to protect the site of pathogen entry as well as actin circulation. Easy administration without needles combined with lower costs for mass immunizations are advantages of mucosal vaccine administration [35]. Adjuvant formulation and delivery are essential, considering that the organization and function of the mucosal immune system differ significantly from the systemic immune system. The *S.pneumoniae* virulence factors pneumolysin and pneumococcal surface protein A are expressed by almost all *S. pneumoniae* serotypes and have been known to play a crucial role in establishing mucosal colonization and subsequent pathogen spreading [24, 36]. Therefore, these proteins have been considered promising protein-based vaccine candidates and were chosen as model antigens for proof of concept to broaden the spectrum of protection by pneumococcal glycoconjugate vaccines.

Antibodies can control and help to clear infections through toxin neutralization and non-neutralizing immune effector functions in addition to preventing pathogen entry. Immunization with ST3-Ply glycoconjugate triggered the production of protective anti-pneumolysin and anti-carbohydrate antibodies. Neutralizing antibodies against the pore-forming toxin pneumolysin inhibited the hemolysis of human red blood cells by whole bacteria cell lysate and prevented alveolar epithelial cell permeability *in vitro.* The *in vivo* model showed that the bacterial burden in blood and lungs, as well as the disease severity, was reduced by immunization with the ST3-Ply conjugate, when mice were challenged with *S. pneumoniae* ST3. The CPS-specific systemic serum antibodies correlated with the opsonophagocytic killing of pneumococci. Hence, intranasal immunization not only inhibits local bacterial colonization but also provides systemic anti-bacterial protection.

Our study illustrates that conjugation of synthetic oligosaccharide antigens to immunogenic pneumococcal carrier proteins (1) provides local protection against colonization in a serotype-independent fashion; (2) triggers a systemic defense by antibody-mediated opsonophagocytic clearance of the bacteria during the infection and (3) inhibits toxin-related side effects by neutralizing Ply-specific antibodies. Further fine-tuning concerning the best combination of protein and synthetic oligosaccharide antigen is still required. Competition with anti◻carrier antibodies elicited by tetanus and diphtheria vaccinations, carrier protein overload, or carrier◻mediated epitope suppression may influence both T◻ and B◻cell responses to glycoconjugate vaccines. Differences in the extent and persistence of protective antibodies triggered by vaccination with bacterial polysaccharide glycoconjugates have been observed in infants [37, 38]. Polysaccharides conjugated to a synthetic multi- or hybrid protein carrier containing multiple T◻cell epitopes might be capable of binding to different types of MHC class II molecules and therefore result in a faster and more robust immune response to polysaccharides when compared with licensed glycoconjugate vaccines [38]. Chimeric protein carriers, such as a combination of PspA and PspC for glycoconjugate vaccines, may further broaden the vaccine spectrum.

Mouse models are often insufficient for addressing particular research questions [39]. The antibody repertoires displayed by mice differ from other mammals [40], and species-specific variations of the immune response to glycan antigen have been observed [41]. Swine and human immune responses are over 80% similar, while human and mouse parameters overlap by less than 10% [42]. Thus, swine were chosen as a large animal model to evaluate the immunogenicity of semisynthetic ST3-Ply and ST3-CRM197 conjugates. This study provides the first evidence for the immunogenicity of synthetic glycoconjugate vaccines in a swine model. The generated antibodies were able to kill pneumococci and neutralize the toxic effect of pneumolysin *in vitro.* However, the protective activity of the glycoconjugate vaccines in the swine *in vivo* infection model requires further investigation. The immune response to a glycoconjugate depends on antibody-producing B cells and on a relatively small population of antigen-specific T lymphocytes that help B cells as well as control the inflammation and protection in response to the pathogen. Understanding the role of antigen-specific T cells in the host immune response is essential for developing effective T cell-dependent vaccines [43]. Here, we demonstrate the involvement of T cells in response to semisynthetic glycoconjugates as well as the specific carbohydrate T cell subpopulation (T_carbs_).

In summary, the global use of PCVs has significantly decreased the burden of IPD caused by serotypes included in the vaccines. However, serotype replacement leads to the increase in invasive diseases caused by non-vaccine types of pneumococcus. Therefore, the strategy of expanding the valency of PCVs, such as PCV15 [44] and PCV24 [45], is a short-term solution. The urgency for a novel universal vaccine is rising up. The combination of synthetic oligosaccharide antigens resembling the pathogen capsule with an immunogenic pneumococcal carrier protein enhanced the efficacy of glycoconjugates against bacterial pneumonia that may be attractive candidates for the development of next-generation vaccines.

## Funding

We gratefully acknowledge generous financial support from the Max Planck Society and the German Research Foundation (SFB/TR 84 “Innate Immunity of the Lung,” C6, C8, C9; to. P.H.S., M.W. and L.E.S.) and by the Federal Excellence Initiative of Mecklenburg Western Pomerania and European Social Fund (ESF) Grant KoInfekt (ESF_14-BM-A55-0001_16 to Sv.H.). We also appreciate the support of Zentrum für Infektionsbiologie und Immunität (ZIBI) Graduate School and International Max Planck Research School for Infectious Diseases and Immunology program (IMPRS-IDI).

## Author contributions

P.K., L.E.S., and P.H.S. designed experiments. P.K. carried out all experiment. P.K., L.Y., K.R. F.V., and T.K. performed a challenge mice study. F.V., T.K., and S.H. provided PspA and Ply protein. A.S. and U.B. conducted pigs immunization. F.E. performed flow cytometry experiment of swine samples. S.H. guided the immunological swine data analyses C.L.P. and S.G.P. synthetized ST3 tetrasaccharide, M.E. synthetized ST2-hexasaccharide. P.M. and M.B. performed conjugation and MALDI characterization. P.K analyzed results and prepared the figures. L.E.S. and P.H.S. conceived of the project. P.K. wrote the initial draft of the manuscript that was edited by L.E.S. and P.H.S. with the input from all authors.

## Competing interests

European Patent Application No. EP 20 170 605.8; Synthetic Streptococcus pneumoniae saccharide conjugates to conserved membrane protein filed by the inventors P.H.S., L.E.S., P.K. and S.V.H. Glycoconjugates containing the synthetic glycan structures of Streptococcus pneumoniae serotypes serotype 2 and 3 are included in patent “Pneumococcal oligosaccharide-protein conjugate composition” no. EP 16 179 133.0 filed by the inventors P.H.S. and P.K.

