## Supplemental Methods for "A semisynthetic glycoconjugate provides expanded cross-serotype protection against *Streptococcus pneumoniae*"

**Materials and Methods**

Conjugation of synthetic oligosaccharides to carrier proteins

Synthetic oligosaccharide *S. pneumoniae* serotype 3 tetrasaccharide (ST3) (Parameswarappa, Reppe et al. 2016) and serotype 2 hexasaccharide (ST2) (Emmadi, Khan et al. 2017) were conjugated to carrier proteins CRM197 (EirGenix, Inc., Taiwan) as well as purified Pneumolysin PlyW433E and PspA using the standardized protocol described previously (Parameswarappa, Reppe et al. 2016, Emmadi, Khan et al. 2017).

Mouse immunization

6-8 weeks-old female C57BL/6N mice (Charles River, Germany) were subcutaneously (s.c.) immunized according to the prime + boost + boost schedule with conjugates corresponding to a dose of 0.4 µg ST3-tetrasaccharide with 0.125 µg of Aluminium hydroxide (Alhydrogen, Denmark) in a total volume of 100 µL PBS. The Prevnar13^®^ group was immunized with 100 µL of commercial vaccine and negative control with PBS.

For the colonization study, mice were immunized intranasally in two-week intervals according to the prime + boost + boost regime with ST2-hexasaccharide-Ply/PspA and ST3-tetrasaccharide-Ply/PspA equal to 2.5 µg of proteins, corresponding to 0.4 - 0.5 µg synthetic oligosaccharide, and adjuvanted with 4 µg of cholera toxin B subunit (CTB; Sigma-Aldrich, United States) in a total volume of 10 µL. PBS and ST3-CRM197 with CTB was administered as a negative control.

S. pneumoniae serotype 3 (ST3) challenge model

*S. pneumoniae* serotype 3 (PN36) were grown in Todd Hewitt Broth with 0.5% (w/v) yeast extract. Cultures were maintained at 37 °C/5% CO_2_ to mid-log growth phase (OD_600_ = 0.3–0.4) and harvested by centrifugation (3100 rpm/10 min). Mice were infected intranasally with 5 x 10^6^ CFU. Disease severity was evaluated at 12-hour intervals and mice were sacrificed 48h after infection. Serial dilutions of blood and lung homogenates were plated on Columbia agar plates with 5% sheep blood to estimate number of CFU.

Intranasal infection of mice with S. pneumoniae serotype 2 (ST2) – colonization model

*S. pneumoniae* serotype 2 (D39) was cultured in liquid THY medium supplemented with 10% heat-inactivated FCS until mid-log growth phase (OD_600nm_ 0.35). Mice were intranasally infected with 3.5 × 10^6^ CFU D39 in 5 µL of PBS. Three days post-infection, mice were euthanized, followed by a collection of nasal tissue and blood.

Swine immunization

Six-week-old, 20 kg female swine (*n* = 30, German landrace *Sus scrofa*) were immunized i.m. according to the prime + boost + boost regime with ST3-CRM197/Ply conjugates and controls, such as Prevnar13^®^, Alum and PBS. A dose of conjugate equal to 2.2 µg of ST3-tetrasaccharide adsorbed onto 125 µg of Aluminium Hydroxide was used.

Glycan array serum screening

Individual synthetic oligosaccharide fragments of *S. pneumoniae* CPS, native CPS, and related proteins were printed on glycan array as previously described (Broecker and Seeberger 2017). Directly before the assay, slides were blocked with the blocking buffer (1% (w/v) BSA in PBS), and incubated with 20-40 μL of the serum samples for 1h at 37 °C. The microarray was washed three times with washing buffer (PBS + 0.1% Tween-20) and the fluorescently labeled secondary antibodies were added.

Enzyme-Linked Immunosorbent Assay - ELISA

ELISA was performed as previously described using high-binding 96-well polystyrene microtiter plates (Corning, USA) coated with different CPSs (SSI Diagnostica, Kopenhagen) at a concentration 10 μg/mL (50 µL per well) in PBS (overnight incubation at 4°C). The plates were washed three times with PBS + 0.1% Tween-20 and blocked with 1% BSA-PBS at RT for 1 h. After three washing steps with PBS + 0.1% Tween-20, plates were incubated with serial dilutions of serum in duplicate or triplicate for 1 h at 37°C. The plates were washed with PBS + 0.1% Tween-20 and treated with horseradish peroxidase (HRP)-labeled secondary antibody diluted in 1% BSA-PBS followed by incubation for 1 h at 37°C. The plates were washed three times with PBS + 0.1% Tween-20, and the color was developed using HRP substrate 3,3'5,5'-tetramethylbenzidine (TMB substrate; BD Biosciences, San Jose, USA). The reaction was stopped by quenching with 2% H_2_SO_4_. The absorbance was recorded at 450 nm using a standard ELISA plate reader.

In Vitro Opsonophagocytic Killing Assay – OPKA

The *in vitro* opsonophagocytic killing assay was performed as described previously by *Romero-Steiner et al.* (Romero-Steiner, Frasch et al. 2003). Briefly, the effector HL-60 cells (a human origin leukemia cell line) at the concentration of approximately 4 × 10^5^ cells/mL in a complete RPMI cell culture medium (90% RPMI 1640, 10% FCS, 1 mM L-glutamine and penicillin-streptomycin solution; PAN Biotech, Germany) were differentiated with 0.8% dimethylformamide (DMF; 99.8% purity; Fisher Scientific, Fair Lawn, NJ, USA) for 5-6 days at 37 °C in the presence of 5% CO_2_. After differentiation, the cells were harvested by centrifugation (300 × g, 5 min), and then viable cells were counted by using 1% trypan blue exclusion and resuspended in opsonophagocytic buffer (HBSS with Ca^2+^ and Mg^2+^, 0.1% gelatin, and 10% FBS; HyClone) at a density of 1 x 10^7^ cells/mL directly before use.

The frozen stock of *S. pneumoniae* previously grown to mid-log phase (in a growth medium at 37 °C / 5% CO_2_ to log phase OD_600_ = 0.3 – 0.4) was diluted in the opsonophagocytic buffer to a final density of 10^6^ CFU/ml (1000 CFU in 20 µL). Individual or pooled sera samples (10 µL per well) were heat-inactivated (56 °C, 30 min) and aliquoted in round bottom non-treated 96-well plates at four-fold dilution intervals (native serum follow by 1:8 to 1:8192 dilutions) and treated with the bacterial suspensions (20 µL per well) to initiate opsonization (incubation for 15 min at 37 °C). After preopsonization, 10 µL of external complement source (baby rabbit complement, CedarLane, Ontario, Canada) as well as 4×10^5^ differentiated HL-60 cells in a volume of 40 µL were added to each well (phagocyte/bacteria ratio 400:1), and plates were incubated for 45 min at 37 °C in 5% CO_2_ environment (preferably with shaking at 220 rpm). The phagocytic reaction was stopped by putting the plate on ice for 20 min. Viable extracellular pneumococci were determined by plating aliquots (5 µL) from each well on Columbia Agar plates with 5% (v/v) sheep blood (BD, New Jersey, USA) and incubating at 37 °C in 5% CO_2_ for several hours to allow bacteria growing (6-8 h in the case of *S.pneumoniae* serotype 3). Visible colony forming units (CFU) were counted. Negative controls lacking either antibody, HL-60, or complement, as well as standard control WHO 007sp typing serum (Human Anti-Pneumococcal capsule Reference Serum, NIBSC, Herts, UK) were used. The assay was repeated two to three times independently. Percentage killing of bacteria was calculated as CFU reduction relative to negative control wells. Serum dilution responsible for 50% killing of bacteria was estimated through non-linear interpolation of the dilution-killing OPKA data

Red blood cell (RBC) lysis assay

The neutralizing ability of Pneumolysin-specific antibodies was tested using the red blood cell lysis assay. A 96-well non-treated round-bottom plate was used for the test. Serum samples were serially diluted in 1% BSA/PBS to a final volume of 25 μL. The full-length native pneumolysin (1 μg/mL) or whole bacteria cell lysates (described below) were added to each well in a volume of 25 μL. The plate was incubated for 30 min at 37 °C with shaking. Human red blood cells (hRBC) were prepared at 1% concentration in PBS and distributed to each well in a volume of 50 μL. The plate was incubated for 60 min at 37 °C without shaking and centrifuged for 10 min at 1000 rpm. 80 μL of supernatants from each well were transferred to a new 96-well flat-bottom plate. The absorbance was measured at 414 nm using an ELISA plate reader. Control wells containing nonlysed 1% hRBC were used to determine the average maximum absorbance value at A_414_. The hemolytic titer was defined as the reciprocal dilution of serum that corresponded to a 50% decrease in the maximum absorbance value at *A*_414_.

Whole bacteria cell lysate preparation

Bacterial culture (2 mL of *S. pneumoniae* serotype 3, serotype 2 and serotype 8), were cultured in brain heart infusion broth (at 37°C in 5% CO_2_ environment) to log phase (OD_600_ = 0.6). Cells were centrifuged, washed with PBS, and resuspended in 100 μL of cell wall digestion buffer (10 mM Tris pH 7.5, 30% sucrose, 1 x protease inhibitor, 1 mg/mL lysozyme) for 3 h at 37 °C. The protoplasts were additionally lysed by sonication (samples on ice, sonication output control 4, 40W, 5 min). The lysates were centrifuged, and the supernatants were filter sterilized. Samples were filled to 1 mL a volume with F-12 K medium (Ham's F-12K (Kaighn's) Medium, Thermo Fisher Scientific, Massachusetts, USA). The supernatant was plated at blood agar plates to confirm the absence of bacterial colonies

Electric Cell-substrate Impedance Sensing - ECIS

The A549 adenocarcinomic human alveolar basal epithelial cell line (received from Sander Group, Department of Infectious Diseases and Respiratory Medicine at Charité Universitätsmedizin Berlin) was maintained in complete DMEM (Dulbecco Modified Eagle Medium‎, PAN Biotech, Germany) with 20% FCS (PAN Biotech, Germany) at a cell concentration between 6 x 10^3^ and 6 x 10^4^ cells/cm^2^.

The ECIS^®^ Zθ Array station (Applied Biophysics Inc, New York, USA) was used for the assay. 1 x 10^5^ A549 cells in a volume of 400 µL of complete DMEM with 20% FCS were added to each well of 8W10E ECIS Cultureware™ Disposable Electrode Array (Applied Biophysics Inc, New York, USA) and cultured for 24h at 37 °C/5% CO_2_. On the next day, the medium was exchanged to DMEM without FSC (100 µL), and the control resistance was measured for one hour. The whole bacteria cell lysates of *S.pneumoanie* serotype 2, 3, and 8 were pre-incubated for 1h at 37°C on a shaker with serum from mice immunized with ST3-Ply conjugate collected on day 35. Pre-incubated bacteria lysate and serum samples were added to the respective wells. The assay was performed overnight. The electrode system measured the resulting voltage (*V*) across the electrodes. The impedance (*Z*) was given by the AC equivalent of Ohm's law: *Z=V/I*, where I was corresponding to a current. The pure resistive (*R*), as well as the capacitive portions (*C*) of the impedance, were also recorded.

Ag-reactive T cell staining of pigs by flow cytometry

Blood was collected into Li-heparin tubes, diluted twice in NaCl, and carefully overlaid on 3 mL of Pancoll, without mixing phases. Samples were centrifuged at 800 x g for 20 minutes, RT and the lymphocytes, together with monocytes and platelets, were harvested. The separated cells were washed twice with RPMI-1640, diluted to the concentration of 12.5 x 106 cells per mL in supplemented media (IMDM (PAN Biotec) supplemented with 10% FCS, 1% P/S, and L-Glu (all purchased from PAN Biotec)). Cells were seeded in the volume of 200 μL PBMC/well in triplicates and stimulated accordingly to the primary immunization either with CRM197 (40 μg/mL), native pneumolysin (40 μg/mL), ST3-CRM197 conjugate (40 μg/mL protein) or ST3-Ply conjugate (40 μg/mL protein). After 2 h, Brefeldin A (ThermoFisher) was added in 1:1000 dilution and incubated for another 4h. The following antibody staining panel was used: dead cell exclusion with fluorophore eFluor 506 Fixable and viability dye (1:500, ThermoFisher, Waltham, USA); anti-human CD14 VioGreen™ 1:50 (clone TÜK-14, Miltenyi Biotech); anti-pig CD3e - PerCp-Cy5.5 1:200 (clone BB23-8E6-8C8, BD Biosciences), anti-pig CD8a – FITC 1:100 (clone 76-2-11, BD Biosciences), anti-pig CD4 – Alexa647 (clone 74-12-4, BD Biosciences), anti-human CD154 - Pe-Vio770 1:50 (clone 5C8, Miltenyi Biotech), anti-pig IFNγ – PE 1:400 (clone P2G10, BD Biosciences), anti-human TNFα – PB 1:100 (Mab11, Biolegend, San Diego, USA). For intracellular staining, Foxp3-fixation and permeabilization kit (ThermoFisher Scientific) was used. Cells were acquired on a FACS CantoII (BD Biosciences), and fcs data was evaluated using Flowjo version 9 (Treestar).

Broecker, F. and P. H. Seeberger (2017). "Synthetic Glycan Microarrays." Methods Mol Biol **1518**: 227-240.

Emmadi, M., N. Khan, L. Lykke, K. Reppe, G. P. S, M. P. Lisboa, S. M. Wienhold, M. Witzenrath, C. L. Pereira and P. H. Seeberger (2017). "A Streptococcus pneumoniae Type 2 Oligosaccharide Glycoconjugate Elicits Opsonic Antibodies and Is Protective in an Animal Model of Invasive Pneumococcal Disease." J Am Chem Soc **139**(41): 14783-14791.

Parameswarappa, S. G., K. Reppe, A. Geissner, P. Menova, S. Govindan, A. D. Calow, A. Wahlbrink, M. W. Weishaupt, B. P. Monnanda, R. L. Bell, L. A. Pirofski, N. Suttorp, L. E. Sander, M. Witzenrath, C. L. Pereira, C. Anish and P. H. Seeberger (2016). "A Semi-synthetic Oligosaccharide Conjugate Vaccine Candidate Confers Protection against Streptococcus pneumoniae Serotype 3 Infection." Cell Chem Biol **23**(11): 1407-1416.

Romero-Steiner, S., C. Frasch, N. Concepcion, D. Goldblatt, H. Kayhty, M. Vakevainen, C. Laferriere, D. Wauters, M. H. Nahm, M. F. Schinsky, B. D. Plikaytis and G. M. Carlone (2003). "Multilaboratory evaluation of a viability assay for measurement of opsonophagocytic antibodies specific to the capsular polysaccharides of Streptococcus pneumoniae." Clin Diagn Lab Immunol **10**(6): 1019-1024.
