## Supplemental Fig.1 for "A semisynthetic glycoconjugate provides expanded cross-serotype protection against *Streptococcus pneumoniae*"

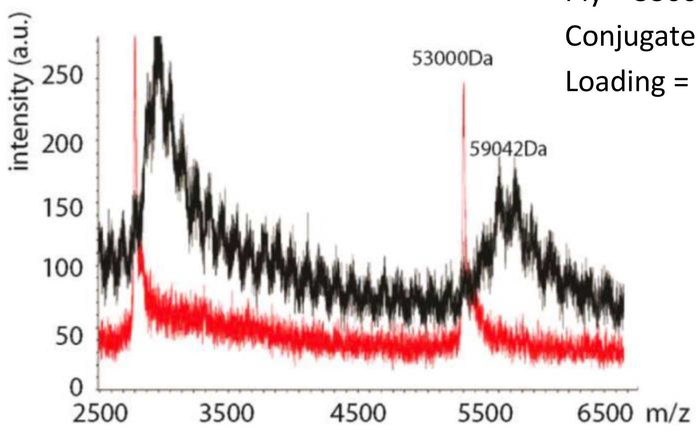


Ply = 53000 Da

Conjugate = 59042 Da

Loading = (59042 – 53000)/864.75 = 6.9 (~7)

**Supplementary Figure 1. Glycoconjugate characterization.** Matrix-assisted laser desorption/ionization (MALDI) analysis was employed to measure the average molecular size of the ST3-Ply conjugate and the pneumolysin carrier protein as a standard.
