## Supplemental Fig.2 for "A semisynthetic glycoconjugate provides expanded cross-serotype protection against *Streptococcus pneumoniae*"

a)

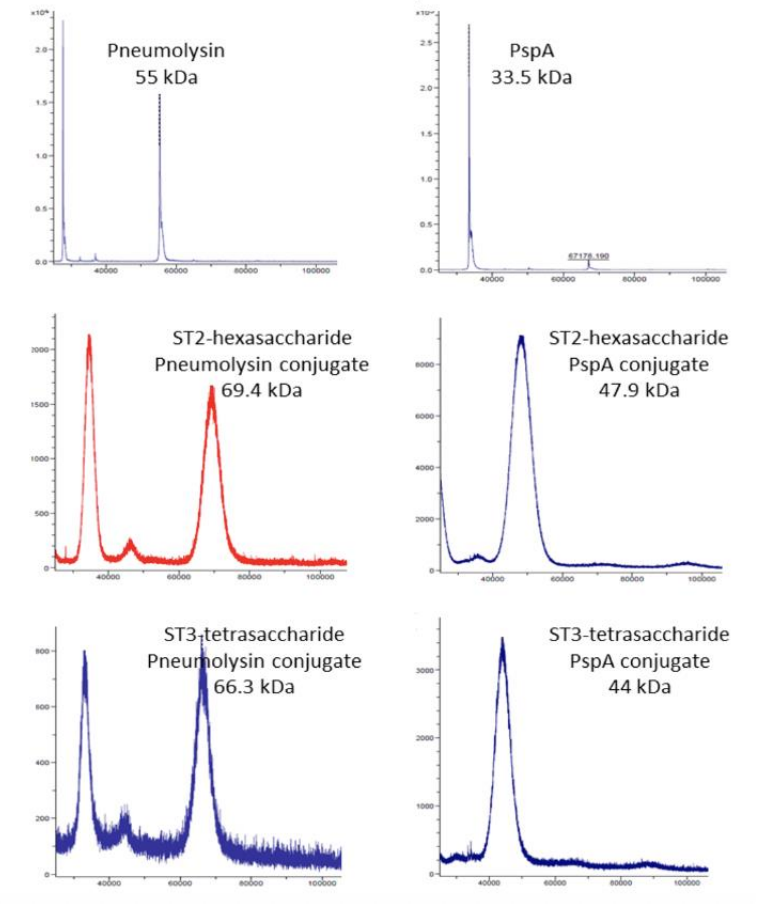


| Construct name | loading |
|---|---|
| SP2-Ply | 12.2 |
| SP2-PspA | 12.6 |
| SP3-Ply | 12.9 |
| SP3-PspA | 12.5 |

**Supplementary Figure 2. Characterization of ST2 and ST3 conjugates.** Matrix-assisted laser desorption/ionization (MALDI) analysis was used to measure the average molecular size of the ST2 and ST3 conjugates. Pneumolysin and PspA proteins were used as a standard.
