## Supplemental Fig.3 for "A semisynthetic glycoconjugate provides expanded cross-serotype protection against *Streptococcus pneumoniae*"

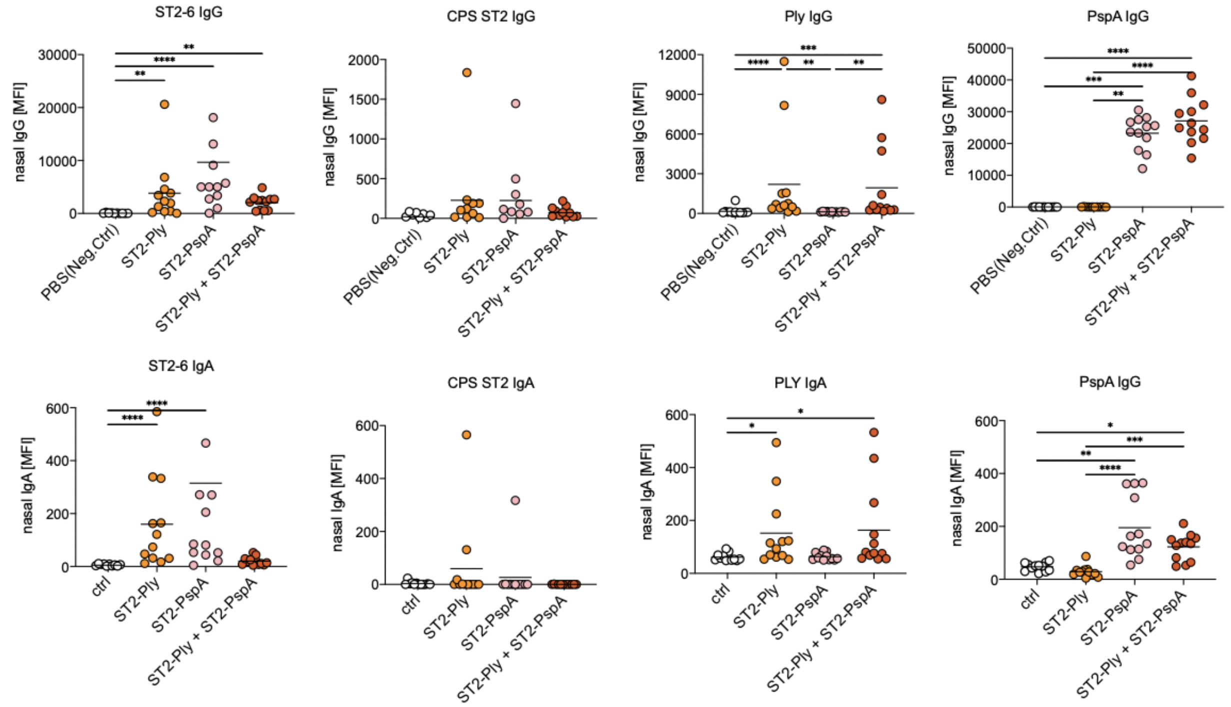


**Supplementary Figure 3.** Evaluation of mucosal Ig**G** and IgA responses on day 3 post intranasal immunization with the indicated vaccines to the indicated antigens (ST2-hexasaccharide (ST2-6); CPS of ST2; Pneumolysin (Ply) and PspA) measured in homogenized nasal tissue by glycan array expressed as MFI. Each dot represents an individual mouse (*n* = 12). Statistical analysis was performed by Tukey's multiple comparisons test; *p < 0.05, **p < 0.01, ****p < 0.001.
